# Mitochondrial ROS1 increases mitochondrial fission and respiration in oral squamous cancer carcinoma

**DOI:** 10.1101/2020.04.13.038844

**Authors:** Yu-Jung Chang, Kuan-Wei Chen, Linyi Chen

## Abstract

Increased *ROS1* oncogene expression has been implicated in the invasiveness of human oral squamous cell carcinoma (OSCC). The cellular distribution of ROS1 has long been assumed at the plasma membrane. However, a previous work reported a differential cellular distribution of mutant ROS1 derived from chromosomal translocation, resulting in increased carcinogenesis. We thus hypothesized that cellular distribution of up-regulated ROS1 in OSCC may correlate with invasiveness. We found that ROS1 can localize to mitochondria in the highly invasive OSCC and identified a mitochondria-targeting signal sequence in ROS1. We also demonstrated that ROS1 targeting to mitochondria is required for mitochondrial fission phenotype in the highly invasive OSCC cells. OSCC cells expressing high levels of ROS1 consumed more oxygen and had increased levels of cellular ATP levels. Our results also revealed that ROS1 regulates mitochondrial biogenesis and cellular metabolic plasticity. Together, these findings demonstrate that ROS1 targeting to mitochondria enhances OSCC invasion through regulating mitochondrial morphogenesis and cellular respiratory.

**Summary Statement:** This study discovers a new role for the ROS1 in mitochondrial fission and metabolic activities.

## INTRODUCTION

Oral cancer is the sixth most common cancer worldwide, and approximately 90% of oral cancers are oral squamous cell carcinomas (OSCCs). Unfortunately, most patients with oral cancer are diagnosed at an advanced stage with neck lymph-node metastasis (Godeny, 2014). The monoclonal antibody therapeutic cetuximab, which targets epidermal growth factor receptor (EGFR), is the most commonly prescribed treatment for advanced OSCC (Loeffler-Ragg et al., 2008), although its clinical efficacy is limited (Bonner et al., 2006, Vermorken et al., 2008) owing to drug resistance or lack of response (Vermorken et al., 2007). Therefore, understanding the molecular mechanisms underlying OSCC metastasis is essential for designing more effective therapeutic approaches.

Receptor tyrosine kinases (RTKs) are synthesized in the endoplasmic reticulum, delivered to the Golgi, and then targeted to the plasma membrane as single-transmembrane proteins where they transduce extracellular signals to orchestrate diverse physiological responses (Geva and Schuldiner, 2014). The *ROS1* oncogene encodes an RTK containing a large N-terminal extracellular domain, a single-transmembrane domain, and an intracellular C-terminal tyrosine kinase domain. We previously showed that upregulation of *ROS1* oncogene leads to oral cancer metastasis (Shih et al., 2017). In addition, genomic rearrangements in *ROS1* have been implicated in cancer progression (Davies and Doebele, 2013). Unlike ROS1 which is assumed to localize at the plasma membrane (Acquaviva et al., 2009), ROS1 fusion proteins derived from chromosomal translocations exhibit differential subcellular localizations and their oncogenic potential can vary. For example, in glioblastoma, the FIG-ROS1 oncoprotein, containing a Golgi-targeting signal, targets to the Golgi rather than the plasma membrane resulting in oncogenic transformation (Charest et al., 2003). Similarly, the endosome-localized fusion proteins SDC4-ROS1 and SLC34A2-ROS1 and endoplasmic reticulum-localized fusion protein CD74-ROS1 differentially activate mitogen-activated protein kinase (MAPK) (Neel et al., 2019). A study has shown that ROS1 localizes to the cytoplasm in majority of OSCCs, whereas it localizes predominantly to the nucleus in adjacent dysplastic epithelial tissues (Cheng et al., 2015). Although this study did not explain whether ROS1 detected by immunofluorescence staining was wild type or had modification, their results suggest that the subcellular distribution of ROS1 may be involved in the pathogenesis of OSCC. To address this issue, we examined the subcellular localization of ROS1 and the results provide new insight into the role of ROS1 in regulating OSCC invasiveness.

## RESULTS

### ROS1 is present in mitochondria

To determine the subcellular localization of full-length ROS1, we first predicted its subcellular distribution using the PSORT-II server. ROS1 was predicted to localize not only to the plasma membrane but also to other organelles such as the endoplasmic reticulum, nucleus, and mitochondria (Fig. 1A). As ROS1 is an RTK, it is not surprising that ROS1 could transit the endoplasmic reticulum during its normal translocation to the plasma membrane. In addition, no obvious plasma membrane or nuclear localization of ROS1 was shown (Fig. 1B and 1C). The fact that Cheng and colleagues reported higher expression of ROS1 in the cytoplasm of OSCC tissues than that in the adjacent dysplastic epithelial tissues suggests that the subcellular localization of ROS1 may be involved in disease progression (Cheng et al., 2015). We also observed that ROS1 and mitochondria co-localized in the highly invasive OSCC cell line C9-IV2 that expresses high levels of ROS1 (Fig. 1B), and these cells are more invasive than the parental C9 cells (Shih et al., 2017). The color map of ROS1 and mitochondrial co-localization was generated using the normalized mean deviation product (nMDP) value, and the index of correlation (Icorr) was calculated using the Colocalization Colormap plugin of ImageJ (Fig. 1B, right bottom panel). Icorr indicated significant co-localization of ROS1 with mitochondria (Icorr = 0.717). We performed the same experiment using COS7 cells transfected with a ROS1-myc construct. Confocal imaging revealed that ROS1-myc co-localized with the mitochondrial outer membrane protein TOM20 (Icorr = 0.811; Fig. 1C). We next used biochemical fractionation of the highly invasive OSCC cell line OC3-IV2 to confirm the subcellular location of ROS1, revealing that ROS1 was indeed present in both the mitochondrial and cytosolic fractions (Fig. 1D). Similarly, the localization of ROS1-myc in 293T cells transfected with a ROS1-myc construct was examined using biochemical fractionation; ROS1-myc was distributed to both mitochondria and cytosol (Fig. 1E). To examine the sub-mitochondrial locale of ROS1, the mitochondrial fractions collected from OC3-IV2 cells were subjected to digestion with proteinase K. ROS1 and TOM20 were digested similarly by proteinase K at 0.5 to 2 μg/ml, whereas the mitochondrial inner membrane protein TIM23 remained protected (Fig. 1F), suggesting that ROS1 is likely a mitochondrial outer membrane protein.

**Fig. 1.**
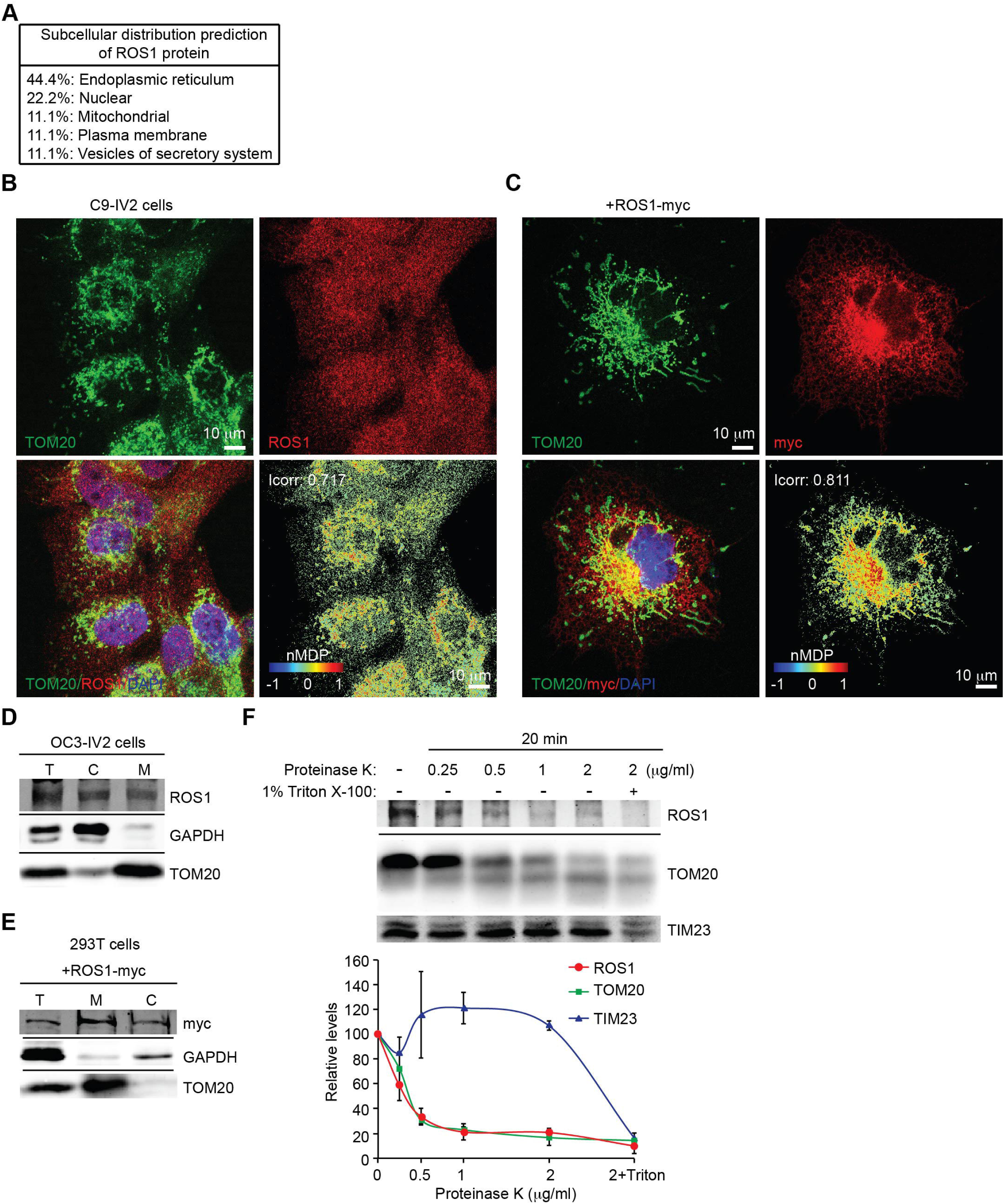
Mitochondrial localization of ROS1. (A) Predicting the subcellular distribution of ROS1 using PSORT-II analysis (http://psort.hgc.jp/form2.html). (B) C9-IV2 cells were subjected to immunostaining for endogenous ROS1 (red) and TOM20 (mitochondria, green) together with DAPI (nucleus, blue). Colocalization Colormap images of ROS1 and TOM20 (bottom right panel) was shown in the normalized mean deviation product (nMDP) value. nMDP value: range 0 to 1, colocalization (warm colors); range −1 to 0, non-colocalization (cold colors). Icorr, index of correlation. (C) COS7 cells transfected with ROS1-myc were subjected to immunostaining for myc (red) and TOM20 (green) together with DAPI (blue). Colocalization Colormap images are shown in the bottom right panel. (D) Subcellular fractionation of OC3-IV2 cells followed by immunoblotting with anti-ROS1, anti-GAPDH, and anti-TOM20. GAPDH and TOM20 served as markers of the cytoplasm and mitochondria, respectively. T: total cell lysate, C: cytoplasmic fraction, M: mitochondrial fraction. (E) Mitochondria and cytosol were subjected to subcellular fractionation of 293T cells transfected with ROS1-myc. Proteins were analyzed by immunoblotting for ROS1, GAPDH, and TOM20. (F) Mitochondria isolated from OC3-IV2 cells were treated with various concentrations of proteinase K for 20 min. Isolated mitochondria that were treated with both proteinase K and Triton X-100 (1%) served as the positive control. Proteins were analyzed by immunoblotting for ROS1, TOM20, and TIM23. The bottom curve plot shows the relative protein level for ROS1, TOM20 and TIM23 after treatment with different concentrations of proteinase K. Proteins were quantified and normalized to the amounts in the no-treatment control. Data from three independent experiments are presented as mean ± SEM. Scale bar: 10 μm in (B-C).

### ROS1 contains a novel mitochondria-targeting signal

Based on the immunostaining and biochemical fractionation results for ROS1 distribution in mitochondria, we next determined which region of ROS1 is responsible for mitochondria targeting. Mitochondrial proteins that are encoded by nuclear genes contain both a moderately hydrophobic transmembrane (T) domain with positively charged residues (P) that flank the T domain and target the proteins to mitochondria (Lee et al., 2014). In addition to three well-defined regions of ROS1, namely the N-terminal extracellular region, T domain, and C-terminal kinase domain, we discovered that the T domain is hydrophobic with a basic positively charged flanking region (residues 1883–1894, defined as the P region; Fig. 2A, left panel). The PSORT WWW prediction server revealed that the TP segment of ROS1 has a 30.4% possibility of being responsible for ROS1 localization to mitochondria. To determine whether the TP segment of ROS1 constitutes a functional mitochondria-targeting signals, we attached GFP to the C-terminus of the TPC region [ROS1(TPC)-GFP], T alone [ROS1(T)-GFP], TP segment [ROS1(TP)-GFP], the PC region [ROS1(PC)-GFP] or C-terminal kinase domain alone [ROS1(C)-GFP] (Fig. 2A, right panel). 293T cells were transfected with these mutant constructs individually, and their expression was confirmed by immunoblotting (Fig. 2B).

**Fig. 2.**
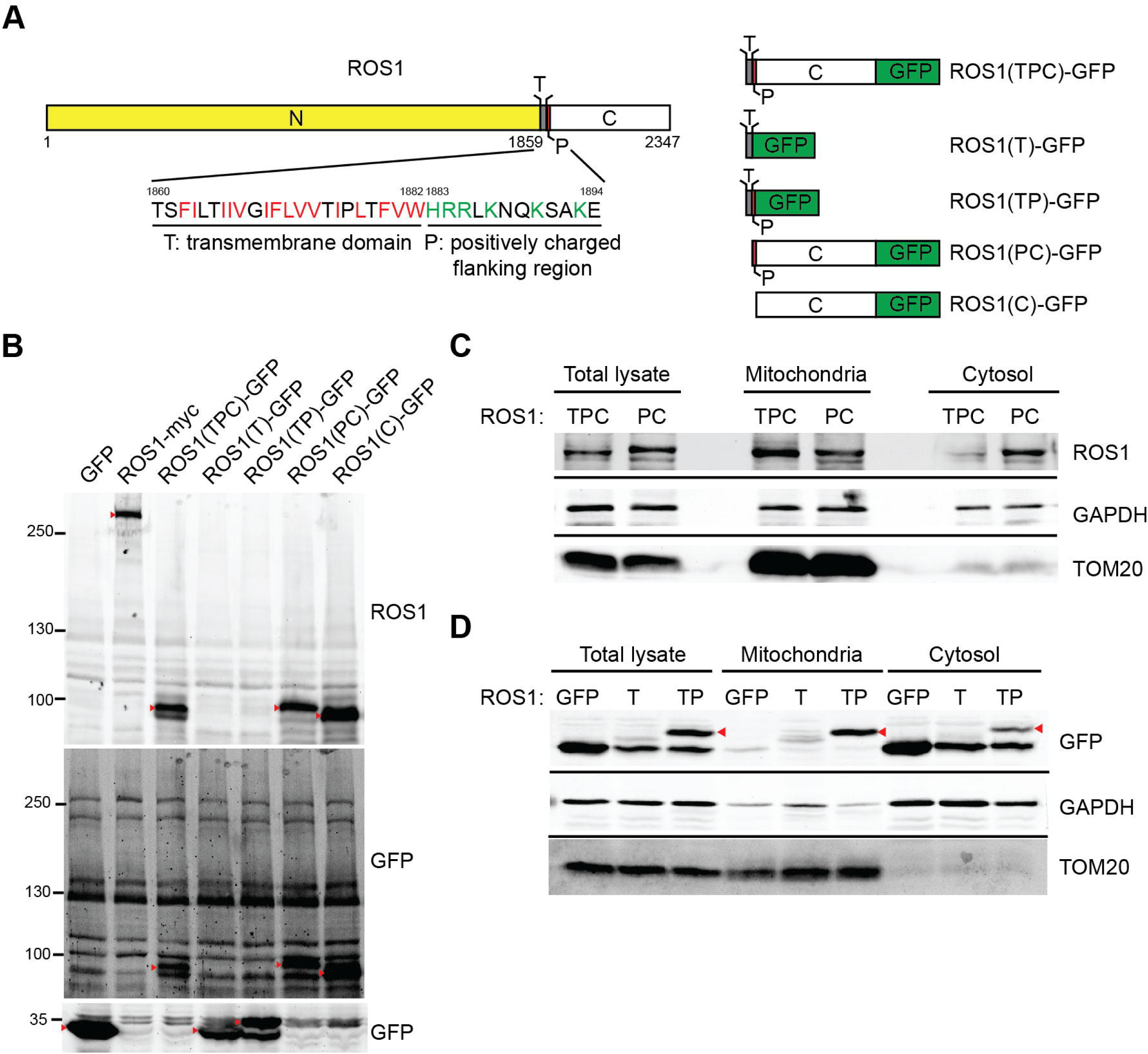
Identification of a mitochondrial targeting sequence in ROS1. (A) Schematic representation of ROS1 mutants. N, N-terminal region; T, transmembrane domain (hydrophobic residues, red); P, positively charged flanking region (positively charged residues, green); C, C-terminal kinase domain. (B) 293T cells were transiently transfected with GFP, ROS1-myc, ROS1(TPC)-GFP, ROS1(T)-GFP, ROS1(TP)-GFP, ROS1(PC)-GFP, or ROS1(C)-GFP. Cell lysates were resolved via SDS-PAGE followed by immunoblotting with anti-ROS1 and anti-GFP. Red arrowheads indicate the proteins of interest. (C) 293T cells were transiently transfected with ROS1(TPC)-GFP or ROS1(PC)-GFP, fractionated, and immunoblotted with anti-ROS1, anti-GAPDH, and anti-TOM20. (D) Mitochondrial and cytosol fractions of 293T cells transfected with GFP, ROS1(T)-GFP or ROS1(TP)-GFP were subjected to subcellular fractionation. Proteins were analyzed by immunoblotting for ROS1, GAPDH, and TOM20.

We first investigated whether the hydrophobic T domain is required for mitochondrial delivery, 293T cells were transfected with the ROS1(TPC)-GFP or ROS1(PC)-GFP constructs, and the localization of the expressed ROS1(TPC) or ROS1(PC) was determined by subcellular fractionation. A significant amount of ROS1(TPC) was present in the mitochondrial fractions (Fig. 2C), whereas ROS1(PC) was primarily detected in the cytosol. This finding suggests that the hydrophobic T domain of ROS1 is important for mitochondrial targeting. To further examine whether the flanking P region of the T domain contributes to ROS1 localization to the mitochondria, we analyzed the localization of GFP, ROS1(T)-GFP, and ROS1(TP)-GFP. Fig. 2D showed that the TP region of ROS1 was able to bring GFP protein into the mitochondria, whereas the T domain alone cannot, suggesting that the P region is required for ROS1 localization to mitochondria. These results indicated that the TP region of ROS1 constitutes a functional mitochondria-targeting signal.

### Mitochondria are more fragmented in highly invasive oral cancer cells

Increased expression of the *ROS1* oncogene promotes OSCC metastasis (Shih et al., 2017), and cancer metastasis is also linked to dysregulated mitochondrial morphogenesis (Zhao et al., 2013, Ma et al., 2019). In addition, ROS1 localizes to mitochondria in the highly invasive OSCC cells (Fig. 1). These data prompted us to investigate whether mitochondrial ROS1 could enhance OSCC invasiveness by affecting mitochondrial morphogenesis. To address this issue, we compared the mitochondrial morphology of isogenic pairs of the highly invasive OSCC cell lines OC3-IV2 and C9-IV2 to that of their respective parental lines OC3 and C9 cells. Immunofluorescence staining and confocal microscopy for TOM20 were used to monitor mitochondrial morphology. Mitochondria were more elongated and tubular in OC3 cells, whereas they were highly fragmented in OC3-IV2 cells and some of the OC3-IV2 mitochondria were of intermediate length (Fig. 3A and 3B). Similarly, C9-IV2 cells contained more fragmented mitochondria than C9 cells (Fig. 3C and 3D). Here, we linked mitochondrial ROS1, fragmented mitochondria and OSCC invasiveness.

**Fig. 3.**
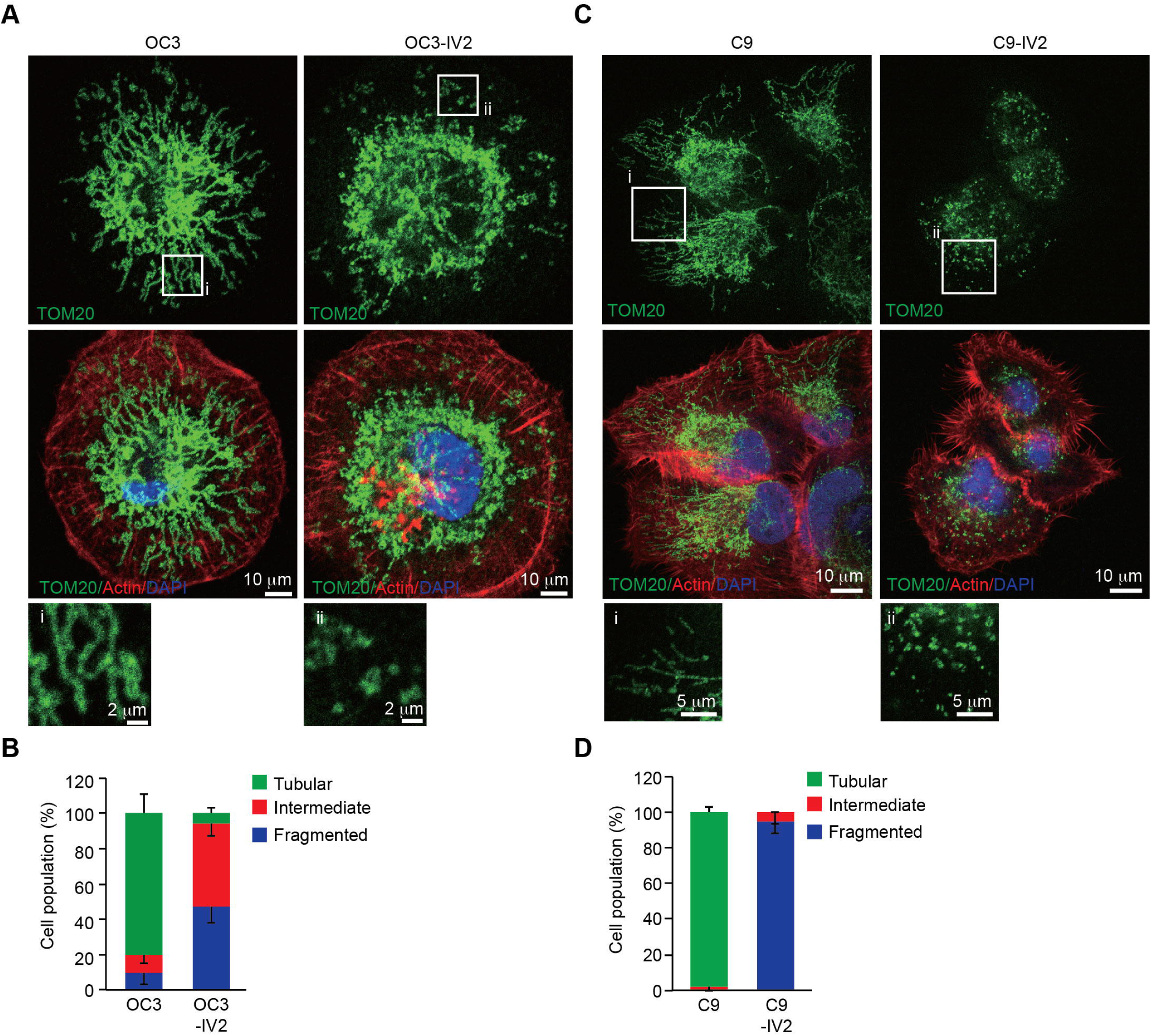
Mitochondria are fragmented in the highly invasive OSCC cells. (A-D) OC3 and OC3-IV2 (A) or C9 and C9-IV2 cells (C) were subjected to immunostaining using anti-TOM20 (green), rhodamine phalloidin (actin, red) and DAPI (nucleus, blue). Enlarged panels from the boxed area are shown in the bottom panel. (B and D) Assessment of mitochondrial morphology in cell lines OC3 (n=68 cells), OC3-IV2 (n=120 cells), C9 (n=22 cells), and C9-IV2 (n=30 cells) in (A and C). Data from at least three independent experiments are presented as mean ± SEM. Scale bar: 10 μm (A and C); 2 μm (A) and 5 μm (C) for enlarged images.

### ROS1 expression induces mitochondrial morphogenesis in OSCC cells

Inhibition of ROS1 activity results in reduced cell proliferation, migration, and invasion of OSCC cells (Shih et al., 2017), so we investigated whether suppression of ROS1 activity could block mitochondrial fission. We treated C9-IV2 cells with DMSO or foretinib (ROS1 inhibitor) and assessed mitochondrial morphology. Foretinib dramatically decreased mitochondrial fission in C9-IV2 cells compared to treatment with DMSO (Fig. 4A and 4B). To determine whether mitochondrial fragmentation of highly invasive OSCC cells occurred in an ROS1-dependent manner, the morphology of mitochondria in OC3-IV2 cells stably expressing a scrambled shRNA (OC3-IV2-Scr) or ROS1-specific shRNA (OC3-IV2-shROS1) was investigated by imaging. In OC3-IV2 cells, ROS1 knockdown reversed the fragmented phenotype to produce elongated mitochondria (Fig. 5A and 5B). In addition, overexpression of ROS1 caused the fused mitochondrial morphology of C9 cells to become fragmented (Fig. 5C and 5D). These data indicated that increased ROS1 and its activity were sufficient to alter mitochondrial morphology.

**Fig. 4.**
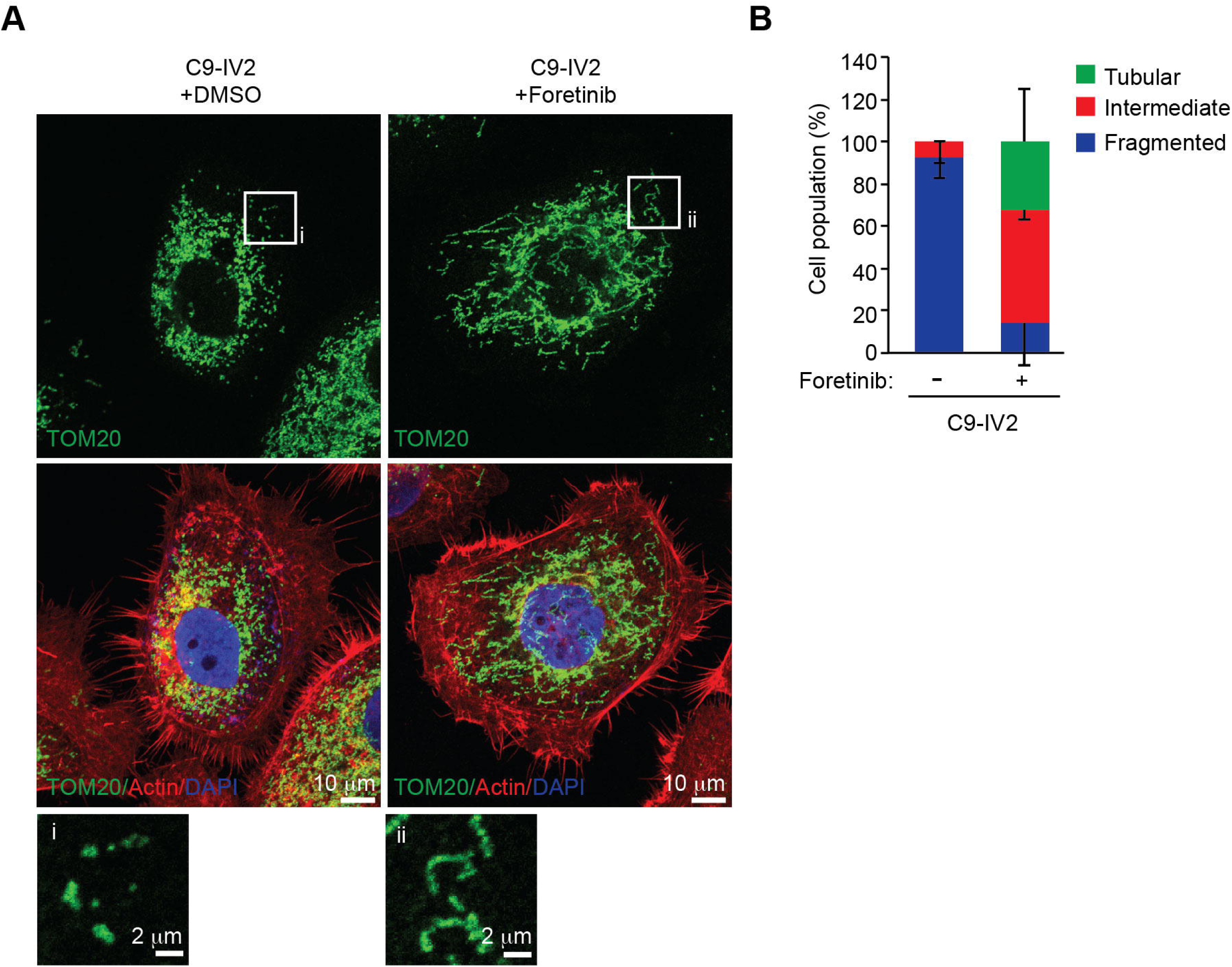
Inhibition of ROS1 signaling leads to mitochondrial fusion. (A) Assessment of mitochondrial morphologies in C9-IV2 cells treated for 24 h with DMSO or 500 nM foretinib (ROS1 inhibitor). Green, TOM20; red, actin; blue, DAPI. (B) Quantification of mitochondrial morphology from (A). Data from three independent experiments are presented as mean ± SEM. Scale bar: 10 μm (A); 2 μm (enlarged images).

**Fig. 5.**
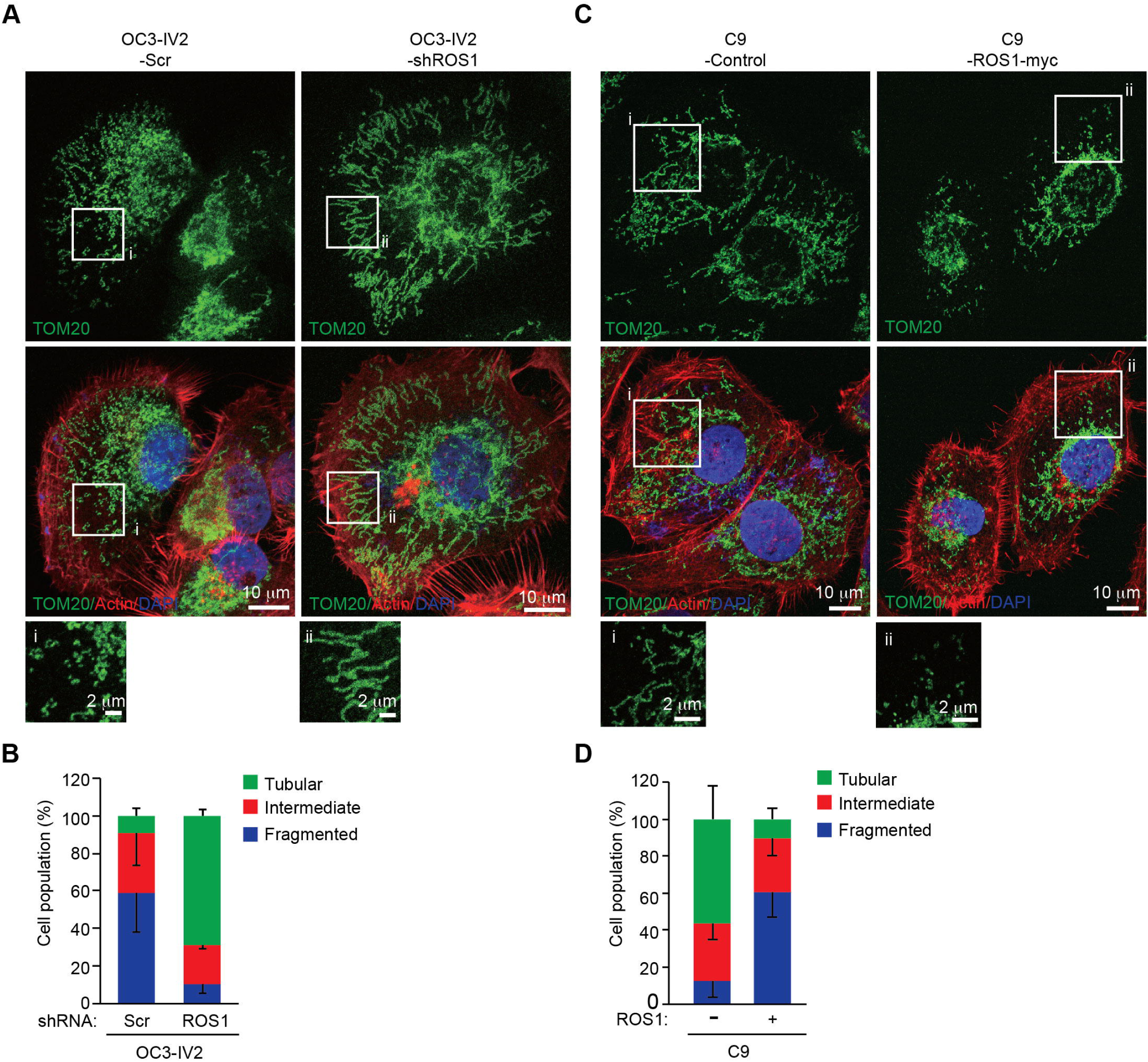
Oncogenic ROS1 is sufficient for mitochondrial fragmentation. (A) OC3-IV2 cells stably expressed an shRNA targeting ROS1 mRNA (OC3-IV2-shROS1) or a scrambled/control shRNA (OC3-IV2-Scr). Mitochondrial morphology was analyzed by immunostaining: green, TOM20; red, actin; blue, DAPI. (B) Quantification of mitochondrial morphology in cell lines OC3-IV2-Scr (n=40 cells) and OC3-IV2-shROS1 (n=35 cells). Data from two independent experiments are presented as mean ± SD. (C) Representative images of the mitochondrial morphologies observed in C9 cells stably transfected with vector only (C9-control) or human ROS1 cDNA (C9-ROS1-myc). Green, TOM20; red, actin; blue, DAPI. (D) Quantification of the mitochondrial morphologies observed in the cells shown in (C). Data in (D) represent the mean ± SEM of three independent experiments with C9-control (n=32 cells) and C9-ROS1-myc (n=51 cells). Scale bar: 10 μm (A and C); 2 μm (enlarged images in A and C).

We next determined whether the mitochondria-targeting signal of ROS1 could affect the morphogenesis of mitochondria. OC3 cells were transfected with the ROS1(TPC)-GFP or ROS1(PC)-GFP constructs and distribution and effect of each protein on mitochondrial morphogenesis were determined by immunofluorescence staining. Confocal images and positive nMDP co-localization values indicated that ROS1(TPC)-GFP localized to mitochondria to a greater extent than ROS1(PC)-GFP (Fig. 6A and 6B). In addition, mitochondria were elongated and tubular in OC3 cells transfected with GFP or ROS1(PC)-GFP, whereas they were mainly fragmented in cells transfected with ROS1(TPC)-GFP (Fig. 6A and 6C). Transfection of COS7 cells with the full-length ROS1 or ROS1-truncated mutant ROS1(TPC) (containing the T domain with the P region) revealed mitochondrial localization of each protein and fragmented mitochondria, whereas this was not the case for COS7 cells transfected with ROS1(PC) (Fig. S1). These results indicated that the newly identified mitochondria-targeting signal is required for mitochondrial delivery of ROS1 and promotes mitochondrial fission.

**Fig. 6.**
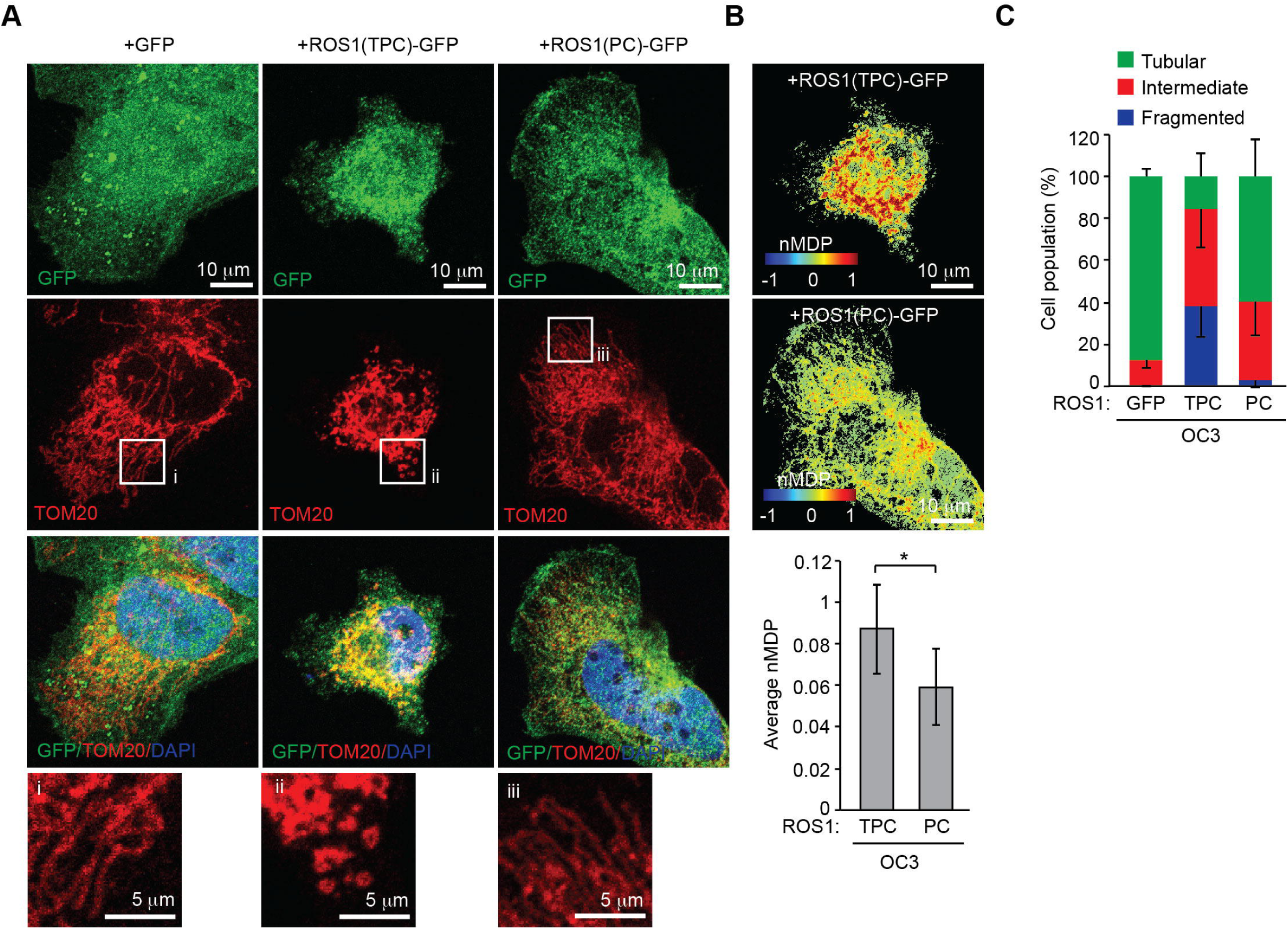
The TP domain of ROS1 is important for ROS1 localization to mitochondria and for ROS1-mediated mitochondrial fragmentation. (A) OC3 cells were transfected with a vector expressing GFP, ROS1(TPC)-GFP, or ROS1(PC)-GFP, and were then immunostained with anti-TOM20 (mitochondria, red) antibody and DAPI (nucleus, blue). (B) Colocalization Colormap images of ROS1 mutants and TOM20 as described in (A). The average of all positive nMDP values represents the abundance of ROS1 mutants in mitochondria. (C) Assessment of mitochondrial morphology in OC3 cells transfected with GFP (n=39 cells), ROS1(TPC)-GFP (n=25 cells), or ROS1(PC)-GFP (n=24 cells) described in (A). Data from at least three independent experiments are presented as mean ± SEM (\**P*<0.05, paired Student’s t-test). Scale bar: 10 μm (A-B); 5 μm (enlarged images in A).

### ROS1 enhances mitochondrial bioenergetics and metabolic plasticity but reduces mitochondrial biogenesis in OSCC cells

The dynamics of mitochondrial morphogenesis correlate with changes in bioenergetics, cell metabolism and motility (Mishra and Chan, 2016). To determine the physiological implications of mitochondrial morphogenesis, we measured the mitochondrial oxygen consumption rate (OCR) and ATP synthesis in OC3 and OC3-IV2 cells. The fraction of basal OCR inhibited by the ATP synthase inhibitor oligomycin was used to estimate the mitochondrial respiration rate required to sustain cellular ATP production. The protonophoric uncoupler FCCP stimulates the mitochondrial electron transport system to function at its maximal capacity, defined as maximal OCR. The spare respiratory capacity is the difference between the basal and maximal OCR in response to a stress. Antimycin A and rotenone inhibit the mitochondrial electron transport system through Complexes I and III to allow measurement of non-mitochondrial respiration processes. The basal OCR was not significantly different between OC3 and OC3-IV2 cells. OC3-IV2 cells, on the other hand, exhibited a 55% enhancement of maximal OCR, a 2-fold elevation in spare respiratory capacity, and a 30% increase in ATP production compared to OC3 cells (Fig. 7A and 7B), reflecting higher energy production in OC3-IV2 cells. This result revealed a link between spare respiratory capacity, ATP production, and cell invasiveness. To examine whether ROS1 directly regulates mitochondrial bioenergetics, we knocked ROS1 in OC3-IV2 cells and determined the effect on mitochondrial OCR. Compared to control OC3-IV2-Scr cells, knockdown of ROS1 reduced basal OCR, maximal OCR, spare respiratory capacity, and ATP production (Fig. 7C and 7D). These results demonstrated that elevated ROS1 enhances mitochondrial bioenergetics.

**Fig. 7.**
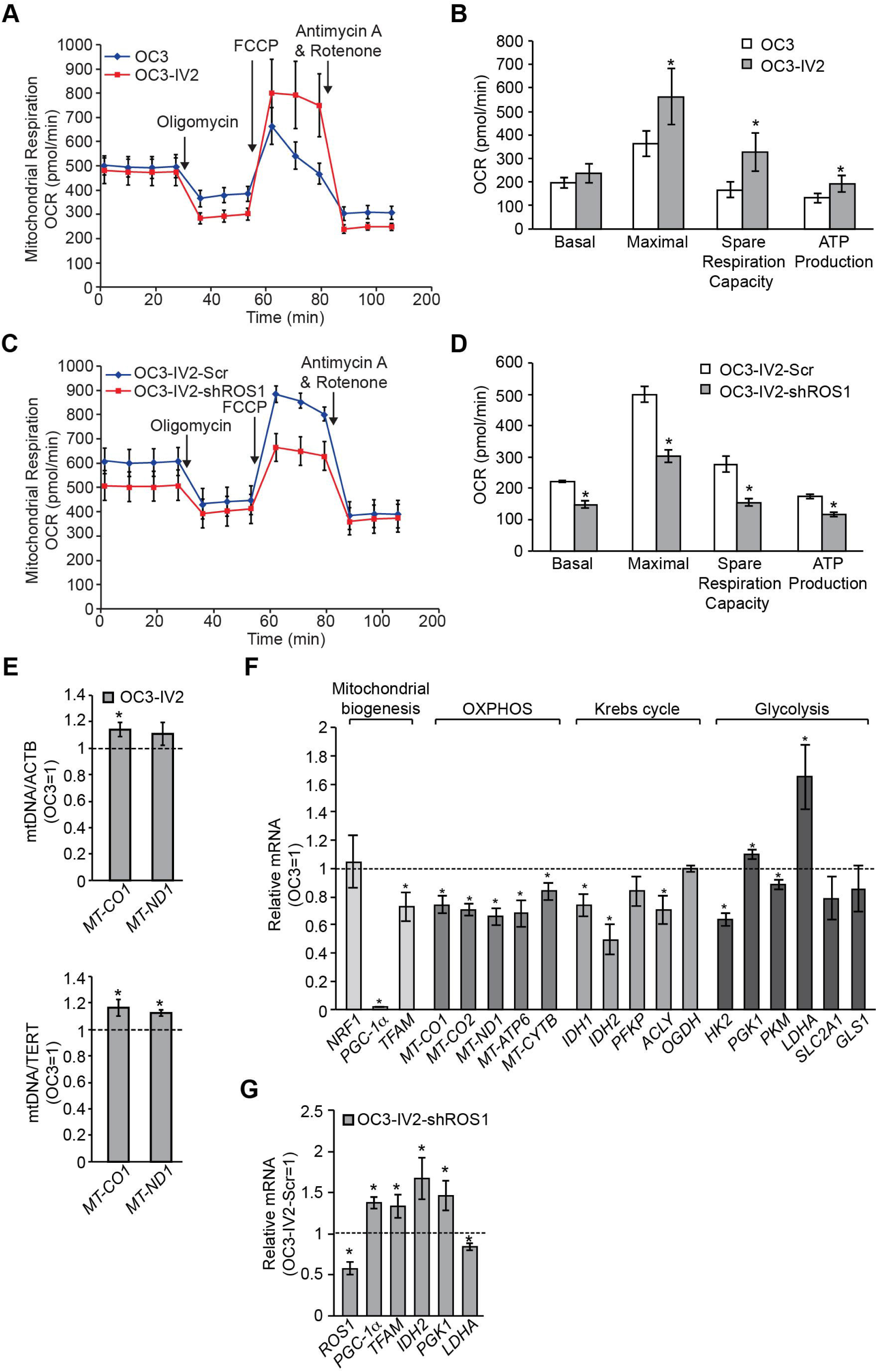
ROS1 affects mitochondrial function in OSCC cells. (A-D) Measurement of oxygen consumption rate (OCR) in OC3 and OC3-IV2 (A) or OC3-IV2-Scr and OC3-IV2-shROS1 cells (C) using the Seahorse XF24 analyzer. (B and D) Average basal OCR, maximal OCR, spare respiration capacity, and ATP production in OC3 and OC3-IV2 (B) or OC3-IV2-Scr and OC3-IV2-shROS1 cells (D) are shown. (E) The relative mtDNA content was measured by Q-PCR. The expression of *MT-CO1* or *MT-ND1* was normalized to nuclear-encoded *ACTB* or *TERT* from OC3 and OC3-IV2 cells. (F) Q-PCR of the mRNA levels of the indicated genes in OC3-IV2 cells normalized to that of OC3 cells. Genes were grouped into functional clusters related to mitochondrial biogenesis, OXPHOS, Krebs cycle, and glycolysis. (G) The mRNA levels of the indicated genes in OC3-IV2-shROS1 cells were normalized to those measured in OC3-IV2-Scr. Data from three independent experiments are presented as mean ± SEM (\**P*<0.05, paired Student’s *t*-test).

It is possible that ROS1-dependent enhancement of mitochondrial bioenergetics may be due to increased mitochondrial biogenesis. To study this issue, we measured the mitochondrial DNA content and expression of genes associated with mitochondrial oxidative phosphorylation (OXPHOS) and mitochondrial biogenesis. OC3-IV2 cells had higher mitochondrial DNA content, whereas the expression of mitochondrial protein-coding genes of the OXPHOS system, namely *MT-CO1, MT-CO2, MT-ND1, MT-ATP6* and *MT-CYTB*, was reduced in OC3-IV2 cells compared to OC3 cells (Fig. 7E-F). Expression of the transcriptional co-factor *PGC-1*α and the master transcription factor *TFAM* for biogenesis was also downregulated in OC3-IV2 cells (Fig. 7F). These data indicated that OXPHOS and ATP production in highly invasive oral cancer cells are concomitant with lower mitochondrial biogenesis. In addition, *PGC-1*α and *TFAM* expressions increased when ROS1 was reduced (Fig. 7G), implicating a direct role for ROS1 in modulating mitochondrial biogenesis.

As efficient mitochondrial OXPHOS has been implicated in cellular metabolism (Cannino et al., 2018), we investigated the effects of the ROS1-induced increase in mitochondrial bioenergetics on expression of genes encoding metabolic enzymes. Compared to OC3 cells, we found that the expression of genes associated with the Krebs cycle and glycolysis, including *IDH1, IDH2, ACLY, HK2* and *PKM*, was reduced in OC3-IV2 cells, whereas *PGK* and *LDHA* were upregulated in OC3-IV2 cells (Fig. 7F). Lactate dehydrogenase A, encoded by *LDHA*, converts pyruvate to lactate and generates ATP, and this pathway has been shown to be an essential source of energy for cancer cells (Kim and Dang, 2006). We also found that ROS1 knockdown led to marked downregulation of *LDHA* (Fig. 7G), suggesting that ROS1 can modulate the expression of *LDHA*. This is consistent with the idea that cancer cells maximize ATP production by both glycolysis and OXPHOS, resulting in the utilization of different nutrients and an increase in metabolic plasticity to maintain tumor survival and metastasis during different stages of disease progression (Lehuede et al., 2016). Together, our findings describe a new role for the *ROS1* oncogene in enhancing mitochondrial bioenergetics and metabolic plasticity to promote OSCC invasiveness.

## DISCUSSION

To our knowledge, this is the first study to demonstrate the mitochondrial distribution of oncogenic ROS1. We also showed that *ROS1* oncogene upregulation in highly invasive OSCC induces mitochondrial fragmentation. Nonetheless, co-immunoprecipitation experiments revealed no interaction between ROS1 and the respective mitochondrial fission and fusion proteins dynamin-related protein 1 (DRP1) and mitofusin 2 (MFN2) (Fig. S2). Although this is a negative result, it remains possible that ROS1, DRP1 or MFN2 may interact transiently at the mitochondrial outer membrane. In vivo proximity ligation assays may allow visualization of their interaction at specific locale. Since ROS1 is known to activate MAPK kinase-extracellular signal-regulated kinase (ERK1/2) and phosphatidylinositol 3-kinase-AKT pathways to promote OSCC cell proliferation and invasion (Shih et al., 2017). We cannot exclude the possibility that ROS1 may regulate mitochondrial morphogenesis through ERK2 and AKT-mediated increases of DRP1 activity (Kashatus et al., 2015, Kim et al., 2016). To identify candidate binding partners for ROS1, we immunoprecipitated ROS1-containing complexes, followed by two-dimensional electrophoresis and mass spectrometry. This approach identified α-enolase, which participates in stabilization of the mitochondrial membrane by binding to an integral mitochondrial membrane protein, VDAC1 (Gao et al., 2014), and protein disulfide isomerase A1, which binds to DRP1, thus reducing the activity of DRP1 and preventing mitochondrial fission (Kim et al., 2018). Thus, ROS1 could potentially interact with DRP1-containing complexes and regulate mitochondrial membrane remodeling.

EGFR is another RTK that undergoes subcellular trafficking from the cell surface to the nucleus, Golgi, or endoplasmic reticulum and influences transcriptional regulation, tumor progression, and drug resistance (Wang and Hung, 2012). EGF-induced endocytosis of EGFR results in the translocation of EGFR to the mitochondria (Demory et al., 2009, Che et al., 2015). Mitochondrial EGFR also induces mitochondrial fission through inhibition of MFN1 and enhances energy production and cell motility to promote metastasis of non-small-cell lung cancer (Che et al., 2015). Mitochondria-localized EGFR associates with c-Src at the mitochondrial inner membrane to phosphorylate cytochrome c oxidase subunit II (COXII), a component of the complex IV of the mitochondrial electron transport system; the interaction leads to inhibition of the COXII activity and to a reduction in ATP level in breast cancer cells (Demory et al., 2009). These two examples reveal multiple functions of RTKs in regulating cancer progress through trafficking to different organelles and potentially being post-translationally modified or cleaved (Chen and Hung, 2015).

What is the physiological implication of the ROS1-induced increase in mitochondrial respiration? One clue from metastatic breast cancer cells is the association between mitochondrial OXPHOS and stemness (Peiris-Pages et al., 2018). We found that the stemness genes *ALDH1, SOX2* and *IL13RA2* were upregulated in OC3-IV2 cells, in which *ROS1* is highly expressed (Fig. S3). However, it remains unknown how ROS1 and increased mitochondrial respiration lead to the upregulation of stemness genes. On the other hand, ROS1 knockdown results in tubular mitochondria and reduced ATP production, which explains its suppression of cell growth, migration and invasion of OSCC cells (Shih et al., 2017).

In summary, our results demonstrate that oncogenic ROS1 is present in mitochondria, governs mitochondrial morphogenesis and confers metabolic plasticity to promote oral cancer invasiveness. Therefore, targeting mitochondrial ROS1 may provide a new therapeutic approach for treating OSCC and prevent OSCC metastasis.

## MATERIALS AND METHODS

### Reagents

Anti-ROS1 (#3266, 1:1000 for immunoblotting, 1:100 for immunofluorescence staining) and anti-DRP1 (#8570, 1:1000 for immunoblotting) antibodies were purchased from Cell Signaling Technology (Danvers, MA, USA). Anti-GAPDH (#G8795, 1:5000 for immunoblotting) and anti-myc tag (#05-724MG, 1:1000 for immunoblotting) antibodies were purchased from Sigma-Aldrich (St Louis, MO, USA). Anti-TOM20 (#sc-11415, 1:1000 for immunoblotting, 1:100 for immunofluorescence staining) and anti-TIM23 (#sc-514463, 1:1000 for immunoblotting) were purchased from Santa Cruz Biotechnology (Santa Cruz, CA, USA). Secondary antibodies conjugated to Alexa Fluor 700 (#A21036), Alexa Fluor 488 (#A11001), Alexa Fluor 555 (#A21428), or Alexa Fluor 647 (#A21235) were purchased from Invitrogen (Carlsbad, CA, USA) and diluted 1:10,000. Rhodamine phalloidin (#A12379) was purchased from Invitrogen and diluted 1:1000 for immunofluorescence staining. IRDye800CW-labeled anti-rabbit secondary antibody (#926-32211, 1:10,000 for immunoblotting) was purchased from LI-COR Biosciences (Lincoln, NE, USA). Anti-GFP tag (#GTX113617, 1:1000 for immunoblotting) and anti-mCherry (#GTX59788, 1:1000 for immunoblotting) antibodies were purchased from GeneTex (Irvine, CA, USA). Foretinib (#A2974) was purchased from ApexBio Technology (Houston, TX, USA).

### Cell culture

COS7 cells and 293T cells were obtained from the American Type Culture Collection, and the human oral cancer cell lines CGHNC9 (C9), C9-IV2, OC3, and OC3-IV2 have been described (Lin et al., 2004, Lu et al., 2012, Huang et al., 2014). OC3 and OC3-IV2 cells were maintained in a 1:1 mixture of Dulbecco’s modified Eagle medium (DMEM; Invitrogen) and keratinocyte serum-free medium (Invitrogen) containing 10% (v/v) fetal bovine serum (Invitrogen), 1% (v/v) L-glutamine (Invitrogen) and 1% (v/v) antibiotic-antimycotic (Invitrogen) and cultured at 37°C and 5% CO_2_. C9, C9-IV2, COS7 cells and 293T cells were maintained in DMEM with 10% (v/v) fetal bovine serum, 1% (v/v) L-glutamine and 1% (v/v) antibiotic-antimycotic and cultured at 37°C under 5% CO_2_ conditions.

### Plasmids

The plasmid encoding human ROS1 (ROS1-myc, #RC220652) was obtained from OriGene Technologies (Rockville, MD, USA). A fragment encoding amino acid resides 1860–1882 (T), 1860–1894 (TP), 1883–2347 (PC) or 1895–2347 (C) of ROS1 was amplified from a ROS1-myc plasmid with PCR using primers for the T domain (forward: 5’-CGGCTAGCATGACAAGTTTCATACT-3’ and reverse: 5’-GACCGGTAGCCAGACAAAG-3’), TP domain(forward: 5’-CGGCTAGCATGACAAGTTTCATACT-3’ and reverse: 5’-GACCGGTAGTTCCTTGGCA-3’), PC domain(forward: 5’-CGGCTAGCATGCATAGAAGATTAAA-3’ and reverse: 5’-GACCGGTAGATCAGACCCATC-3’),or C domain (forward: 5’-CGGCTAGCATGGGGGTGACAGT-3’ and reverse: 5’-GACCGGTAGATCAGACCCATC-3’) and then subcloned into pEGFP-C1 via the NheI/AgeI sites. Myc construct was a gift from Dr. Christin Carter-Su (Rui et al., 1999). mCherry-DRP1 (#49152) and MFN2-YFP (#28010) was obtained from Addgene (Watertown, MA, USA). mCherry-DRP1 was subjected to PCR to introduce cleavage sites at the 5’ and 3’ ends of mCherry (forward: 5’-CGGATCCGGTCGCCACCATGGTGAG-3’ and reverse 5’-CGGACTTGTACAGCTCGTCCA-3’), and the PCR products were subcloned into MFN2-YFP via the BamHI/BsrGI sites to replace YFP and generate the MFN2-mcherry construct.

### Knockdown of ROS1

pLKO.1-TRC2.Void [Scramble (Scr), ASN0000000001] and pLKO.1-shROS1 (TRCN0000219660) were obtained from the National Core Facility at the Institute of Molecular Biology, Genomic Research Center, Academic Sinica, Taiwan. Lentivirus containing pLKO.1-TRC2.Void or pLKO.1-shROS1 was prepared from 293T cells co-transfected with pCMVΔR8.91 and pMD.G along with pLKO.1-TRC2.Void or pLKO.1-shROS1. Medium containing lentivirus was harvested and added into OC3-IV2 cells. OC3-IV2 cells were subjected to selection with 2 μg/ml puromycin for at least 2 weeks.

### Reverse transcription and semi-quantitative real-time PCR (Q-PCR)

RNA was isolated from OSCC cells using TRIzol reagent (Invitrogen) and converted to cDNA using a reverse transcription kit (Applied Biosystems, Foster City, CA, USA). Q-PCR was performed using Power SYBR Green PCR Master Mix and ABI StepOnePlus Real-Time PCR System (Applied Biosystems) with specific primers: ACTB (forward: 5’-AGGATCTTCATGAGGTAGTCAGTCAG-3’ and reverse: 5’-CCACACTGTGCCCATCTACG-3’), TERT (forward: 5’-CAAGCACTTCCTCTACTCCTC-3’ and reverse:5’-TGGAACCCAGAAAGATGGTC-3’), NRF1 (forward: 5’-GGCACTGTCTCACTTATCCAGGTT-3’ and reverse: 5’-CAGCCACGGCAGAATAATTCA-3’), PGC-1α(forward: 5’-AATCAGACCTGACACAACAC-3’ and reverse: 5’-GCACTCCTCAATTTCACCAA-3’), TFAM (forward: 5’-ACTGCGCTCCCCCTTCAG-3’ and reverse: 5’-ACAGATGAAAACCACCTCGGTAA-3’), MT-CO1 (forward: 5’-GGCCTGACTGGCATTGTATT-3’ and reverse:5’-TGGCGTAGGTTTGGTCTAGG-3’), MT-CO2 (forward: 5’-CGATCCCTCCCTTACCATCA-3’ and reverse:5’-CCGTAGTCGGTGTACTCGTAGGT-3’), MT-ND1(forward: 5’-CCAATGATGGCGCGATG-3’ and reverse: 5’-CTTTTTGGACAGGTGGTGTGT-3’), MT-ATP6 (forward: 5’-CCCACTTCTTACCACAAGGC-3’ and reverse:5’-GTAGGTGGCCTGCAGTAATG-3’), MT-CYTB (forward: 5’-CTCCCGTGAGGCCAAATATC-3’ and reverse: 5’-GAATCGTGTGAGGGTGGGAC-3’), IDH1 (forward: 5’-ATGCAAGGAGATGAAATGACACG-3’ and reverse: 5’-GCATCACGATTCTCTATGCCTAA-3’), IDH2 (forward: 5’-TACGGGTCATCTCATCACCA-3’ and reverse: 5’-ACCTCGCAAGAGCAGCC-3’), PFKP (forward: 5’-CGGAGTTCCTGGAGCACCTCTC-3’ and reverse:5’-AAGTACACCTTGGCCCCCACGTA-3’),ACLY(forward: 5’-GAAGGGAGTGACCATCATCG-3’ and reverse: 5’-TTAAAGCACCCAGGCTTGAT-3’), OGDH (forward: 5’-AGATCATCCGTCGGCTGGAGATGG-3’ and reverse: 5’-TCAAACTTCTGCCGGATCCACTG-3’), HK2 (forward: 5’-ATGAGGGGCGGATGTGTATCA-3’ and reverse:5’-GGTTCAGTGAGCCCATGTCAA-3’), PGK1 (forward: 5’-GGGCAAGGATGTTCTGTTCT-3’ and reverse:5’-TCTCCAGCAGGATGACAGAC-3’),PKM (forward: 5’-ATCGTCCTCACCAAGTCTGG-3’ and reverse: 5’-GAAGATGCCACGGTACAGGT-3’), LDHA (forward: 5’-AGCCCGATTCCGTTACCT-3’ and reverse: 5’-CACCAGCAACATTCATTCCA-3’), SLC2A1 (forward: 5’-CCAGCTGCCATTGCCGTT-3’ and reverse: 5’-GACGTAGGGACCACACAGTTGC-3’), GLS1 (forward: 5’-TGGTGGCCTCAGGTGAAAAT-3’ and reverse:5’-CCAAGCTAGGTAACAGACCCTGTTT-3’), ALDH1 (forward: 5’-GTTGTCAAACCAGCAGAGCA-3’ and reverse: 5’-AGGCCCATAACCAGGAACAA-3’), SOX2 (forward: 5’-AGGATAAGTACACGCTGCCC-3’ and reverse: 5’-TTCATGTGCGCGTAACTGTC-3’), IL13RA2 (forward: 5’-AGTTAAACCTTTGCCGCCAG-3’ and reverse: 5’-AGGTCCCAAAGGTATGCTCC-3’) and GAPDH (forward: 5’-CAAGGCTGTGGGCAAGGT-3’ and reverse: 5’-GGAAGGCCATGCCAGTGA-3’). All results of Q-PCR were normalized to *GAPDH*.

### Immunoblotting

Cell were lysed in SDS lysis buffer (240 mM Tris-acetate, pH 7.8, 5 mM EDTA, 1% (w/v) SDS, 0.5% (v/v) glycerol) containing 1 mM sodium orthovanadate (Na_3_VO_4_), 1 mM phenylmethylsulfonyl fluoride, 10 ng/ml leupeptin and 10 ng/ml aprotinin. Protein concentrations were determined by the bicinchoninic acid assay (Santa Cruz Biotechnology), and were separated by SDS-PAGE. The separated proteins were transferred to a membrane and immunoblotted with the primary antibodies indicated in each figure and the appropriate IRDye-conjugated secondary antibody. Immunopositive bands were detected using the Odyssey infrared imaging system (LI-COR Biosciences).

### Immunofluorescence staining, confocal microscopy, mitochondrial morphology, and colocalization analysis

For immunofluorescence staining, cells were fixed in 4% (v/v) paraformaldehyde and permeabilized in 0.1% (v/v) Triton X-100. Cells were then incubated in a blocking buffer containing 1% bovine serum albumin and then with the indicated primary antibodies followed by Alexa Fluor–conjugated secondary antibodies. Nuclei were stained with DAPI, and cells were mounted using Prolong Gold reagent (Invitrogen). Images were acquired using a LSM780 confocal microscope and analyzed with ZEN 2012 software (Zeiss, Jena, Germany). Mitochondrial morphology in cells was categorized into three different groups: tubular, fragmented, and intermediate; Tubular had at least one mitochondrial tubule ≥5 μm in length; intermediate, at least one between 2 and 5 μm, but no longer than 5 μm; fragmented, none ≥2 μm in length. Mitochondrial length was calculated using the Simple Neurite Tracer plugin of ImageJ software. For the colocalization analysis, the Colocalization Colormap plugin of ImageJ was used to generate a color-coded image of colocalization and calculate the nMDP value and Icorr. The nMDP value represents the non-colocalization with cold colors (range −1 to 0) and colocalization with warm colors (range 0 to 1). Icorr indicated the fraction of positively correlated pixels in the image.

### Subcellular fractionation and proteinase K treatment of mitochondria

Mitochondrial and cytoplasmic fractions were extracted using the Mitochondria Isolation kit for Cultured Cells (Thermo Fisher Scientific, #89874, Waltham, MA, USA). Briefly, cells were collected and then incubated with Reagent A for 2 min on ice. Reagent B was added, and the sample was vortexed at maximum speed, followed by incubation for 5 min on ice. Reagent C was added, and the sample was centrifuged at 700 × *g* at 4°C for 10 min. The supernatant was collected and centrifuged at 12,000 × *g* at 4°C for 15 min. The supernatant represented the cytoplasmic fraction, and the pellet represented the mitochondria-enriched fraction. The pellet was washed using Reagent C and then lysed with SDS lysis buffer. For digestion with proteinase K, equivalent weights of the mitochondria-enriched fraction were resuspended in digestion buffer (250 mM sucrose, 0.5 mM EGTA, 0.5 mM EDTA, 3 mM HEPES-NaOH, pH 7.2), and proteinase K (Sigma-Aldrich) was added to a final concentration of 0.25, 0.5, 1 or 2 μg/ml with or without 1% (v/v) Triton X-100, with incubation for 20 min on ice. The reaction was stopped by the addition of 2 mM phenylmethylsulfonyl fluoride, and samples were collected and lysed after centrifugation at 15,000 × *g* at 4°C for 10 min.

### Measurement of OCR

The OCRs were measured using the Seahorse XF24 Analyzer (Taiwan Mitochondrion Applied Technology, http://www.taimito.com/taimito/index.php, Taiwan). Cells were seeded at 1.7 × 10^4^ per well in DMEM/keratinocyte serum-free medium containing 10% (v/v) fetal bovine serum. After 24 h, cells were transferred into assay media and subjected to addition of either 1 μM oligomycin, 1 μM FCCP, 1 μM rotenone, or 1 μM antimycin A, for determination of basal OCR, ATP production, maximal OCR, spare respiratory capacity, respectively.

### Statistical analysis

Statistical analysis of all results was carried out using the paired Student’s *t*-test. All values reflect the mean ± S.E.M. of data obtained from at least three independent experiments if not otherwise specified. Significance was defined as *P* < 0.05.

## Symbols and Abbreviations

COXII: cytochrome c oxidase subunit II;
DRP1: dynamin-related protein 1;
EGFR: epidermal growth factor receptor;
ERK: extracellular signal-regulated kinase;
Icorr: index of correlation;
MFN2: mitofusin 2;
nMDP: normalized mean deviation product;
OCR: oxygen consumption rate;
OSCC: oral squamous cell carcinoma;
OXPHOS: oxidative phosphorylation;
P: positively charged residues;
RTK: receptor tyrosine kinase;
T: transmembrane domain;
TP: transmembrane domain with flanking positively charged residues.

## Author contributions

Conceptualization: LC and YJC; Methodology: YJC and KWC; Validation: YJC; Formal analysis: YJC; Investigation: LC and YJC; Resources: LC and YJC; Data curation: YJC and KWC; Writing - original draft: LC and YJC; Writing - review & editing: LC, YJC, KWC; Visualization: YJC; Supervision: LC; Project administration: LC and YJC; Funding acquisition: LC and YJC.

## Figure legends

**Fig. S1.**
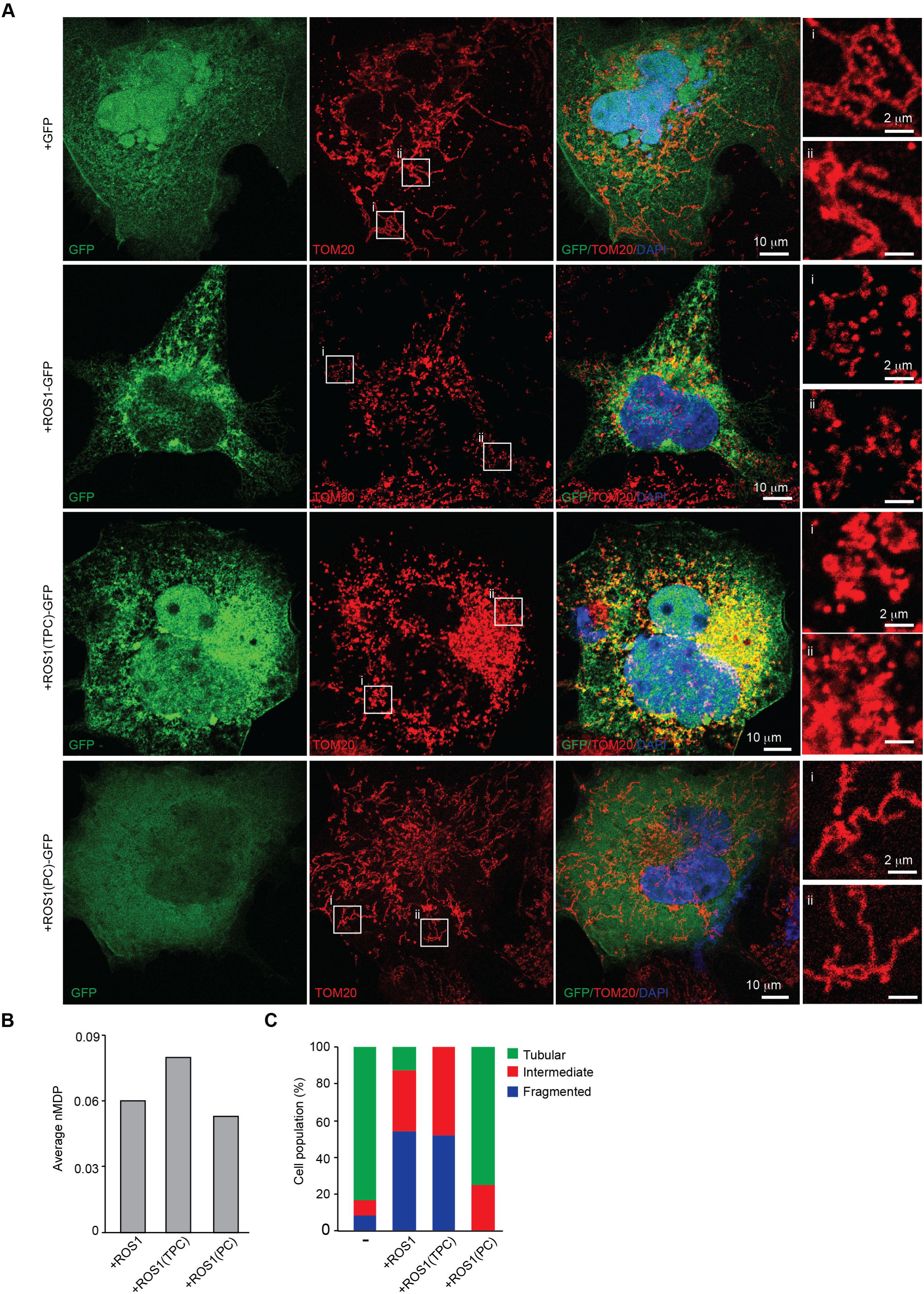
Distribution of ROS1 mutants and their effects on mitochondrial phenotype in COS7 cells. (A) COS7 cells were transfected with a vector expressing GFP, ROS1-GFP, ROS1(TPC)-GFP, or ROS1(PC)-GFP and then immunostained with anti-TOM20 (mitochondria, red) and DAPI (nucleus, blue). (B) Quantification of the average nMDP values. (C) Morphology of mitochondria from COS7 cells transfected with GFP (n=9 cells), ROS1-GFP (n=7 cells), ROS1(TPC)-GFP (n=17 cells) or ROS1(PC)-GFP (n=4 cells) was quantified. Scale bar: 10 μm (A); 2 μm (enlarged images).

**Fig. S2.**
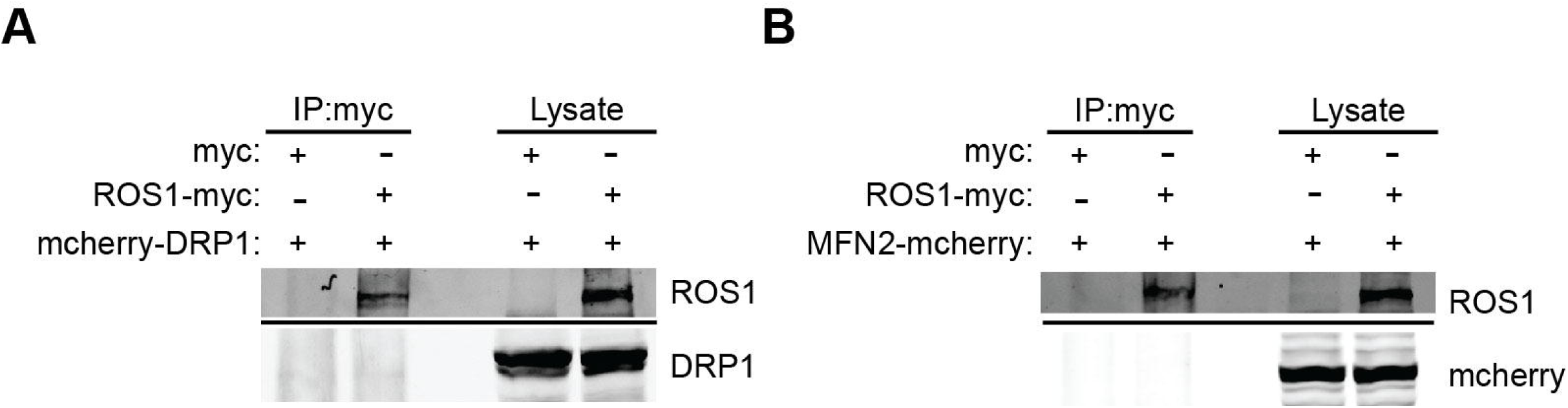
ROS1 oncoprotein does not interact with DRP1 or MFN2. (A-B) Lysates of 293T cells that had been transfected with myc vector or ROS1-myc along with mcherry-DRP1 (A) or MFN2-mcherry (B) were immunoprecipitated using anti-myc and immunoblotted for the indicated proteins.

**Fig. S3.**
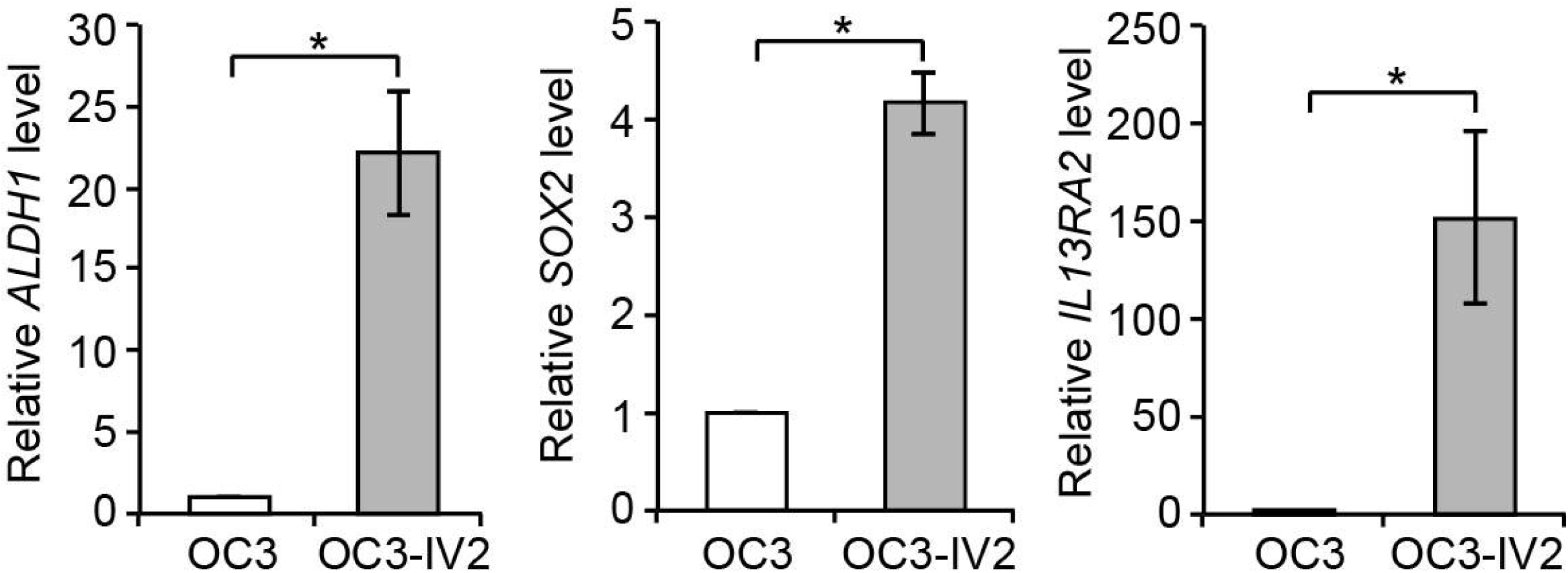
Comparison of stemness gene expression in OSCC cells. Relative expression of *ALDH1, SOX2*, and *IL13RA2* mRNAs in OC3 and OC3-IV2 cells was measured using Q-PCR. Data from three independent experiments are presented as mean ± SEM (\**P*<0.05, paired Student’s *t*-test).

